# Dynamic fusion of structural and functional connectivity via joint connectivity matrix ICA

**DOI:** 10.1101/2025.05.15.653851

**Authors:** Lei Wu, Marlena Duda, Armin Iraji, Vince D. Calhoun

## Abstract

The integration of multimodal MRI images, including functional MRI (fMRI) and diffusion MRI (dMRI), depicts a key advancement in neuroimaging, since it offers a more comprehensive and better understanding of brain function and connections. FMRI captures brain functional activity while dMRI reveals structural connectivity via white matter bundles, each of which provide unique yet complementary insights; however combing these two modalities, particularly at a dynamic level, is challenging due to their drastically different data characteristics. This study introduces a novel framework named “dynamic fusion,” which extends joint component analysis (cmICA) to integrate static structural connectivity (SC) with dynamic functional connectivity (FC). Our approach gauges the relationship between these joint components across various temporal states, aiming to discover both static and dynamic features of brain connectivity. We applied this approach to fMRI and dMRI data from the same set of control subjects, which we had previously studied to estimate joint parcellation and their structural and functional connections using a static model only, and also included a comparable number of individuals with schizophrenia from the same study. Our results reveal that dynamic fusion not only highlights diverse temporal dynamics in FC but, more importantly, also shows how SC patterns differ across dynamic functional states at different time frames, providing new insights into brain organization. Furthermore, it successfully detects joint structural-functional connectivity differences between individuals with schizophrenia and controls, demonstrating its potential for detecting hidden group differences. Overall, this study establishes our dynamic fusion as a powerful tool for integrating structural and dynamic functional connectivity data, enhancing our understanding of brain connectivity and offering new perspectives for studying neurological disorders.

## Introduction

The fusion of functional MRI (fMRI) and diffusion MRI (dMRI) fMRI provides a powerful tool to explore brain function and organization more systematically and organically. Recent emphasis on multimodal neuroimaging, e.g., integrating fMRI and dMRI, has advanced our comprehension of brain communication and its regulation of functional integration and segregation (Chu et al., 2018, Hagmann et al., 2008, Deco et al., 2015). White matter tracts serve as the neuroanatomical pathways connecting functional network regions, facilitating the transmission of interregional information (Honey et al., 2009). Studies of various brain disorders, such as Alzheimer’s disease and dementia, suggest that subtle alterations in white matter connectivity may significantly contribute to the impairment of functional networks associated with cognitive decline (Taylor et al., 2017). Consequently, methods offering enhanced mapping of structural connectivity architecture and functionally linked brain networks through whole-brain modeling (Deco et al., 2015) may play a pivotal role in finding the underlying mechanisms of the brain. However, integrating these two modalities remains challenging and significantly understudied (Calhoun and Sui, 2016, Duda et al., 2024), mainly due to the drastic differences across these two data modalities, including input definitions, characteristics, scales and/or strong assumptions (Wu and Calhoun, 2023).

Among various measurements of fMRI, dynamic functional network connectivity (FNC) has garnered significant attention in recent years, as it enables the capture of moment-to-moment changes in traditional functional connectivity patterns. Dynamic FNCs are typically quantified by pairwise (partial) correlation across two functional areas or networks, at a much shorter temporal scale (20 to 60 seconds) compared to a full scan period (usually above 5 minutes) (Iraji et al., 2021). Such patterns have been demonstrated to highlight dynamic fluctuations across a wide array of conditions during rest or task performance, including brain-mind conditions (Rabinovich et al., 2012, von der Malsburg et al., 2010), vigilance status (Allen et al., 2012, Allen et al., 2018) and free thought (Doucet et al., 2012, Hansen et al., 2015). This has since sparked broad interest in understanding temporal variations in FC across large-scale brain networks/nodes, i.e. the chronnectome (Calhoun et al., 2014, Lurie et al., 2020). Studies of the temporal dynamics of brain organization have shown a high degree of structured temporal variability of the functional connectome, which may be associated with mental status transitions and/or adaptive processes.

To our knowledge, a multimodal connectivity fusion of dMRI and fMRI in the context of dynamic reconfiguration has not yet been thoroughly developed. Here we explore whether brain connectivity organization derived from multimodal fusion with static structural data depends on the dynamic functional state with which it is fused. An initial approach to capture this can be implemented by evaluating the relationship between the identified components across multiple data fusion models. We call this new framework dynamic fusion. Specifically, we applied and extended the joint cmICA approach (Wu and Calhoun, 2023), which provides a data-driven parcellation and automated-linking of SC and FC information simultaneously, to dynamic FC and SC from functional MRI and diffusion-weighted MRI data. This was done by performing multiple fusions and comparing the fusion-static and fusion-dynamic components within each modality. Our prediction was that diffusion data jointly linked to dynamic fMRI data would yield a combination of static (fusion invariant) and dynamic features. That is certain dMRI components would likely be identified regardless of the fMRI data, whereas other dMRI components would, themselves, be dynamic, that is a function of the fMRI state.

The results not only confirmed diverse temporal dynamics in FNC patterns but also revealed that the fusion patterns of FC and SC change over time as well. More intriguingly, different temporally-isolated dynamic functional states are associated with variations in the decomposed SC, i.e., subtle changes in tract pattern vary over time. That is the optimal decomposition of the SC as a function of the temporal state. In addition, the dynamic state links showed differing degrees of variation, with the majority exhibiting more static characteristics while a few links are more dynamic. This finding opens up a new perspective on multimodal imaging. Furthermore, we also showed this technique identified linked structural-functional connectivity changes across individuals with schizophrenia and controls, revealing group differences that would otherwise have been hidden. Interestingly, the group differences tended to be concentrated in the dynamic dMRI states, providing evidence of the importance of optimizing the data decomposition in a way that is jointly informed by multiple modalities.

In the remainder of the manuscript, we first introduce our dynamic fusion framework. Following this, we present cases of dynamically linked patterns in which two distinct fusions were employed to jointly decompose FC and SC, with FC constructed at two different time frames separately. Finally, we analyze the linked connectivity maps from “static” to “dynamic” status, as well as their sensitivity to individuals with schizophrenia versus controls. Overall, our data-driven joint cmICA provides a novel and intriguing approach for integrating SC and dynamic FC. Not only does our approach provide a joint fusion of static SC and FNC, it also provides an innovative and effective tool for functionally aware decomposition of SC.

## Methods

Here, we begin by providing a conceptual overview of our proposed dynamic fusion framework. Next, we present some preliminary results applied to fMRI and dMRI data. Additionally, we evaluate the degree of dynamism exhibited by each component in each modality across fusions and examine its differentiability between schizophrenia and controls.

### 1. A dynamic fusion framework

#### 1.1 Conceptual approach

Figure 1 illustrates the proposed framework. To implement the framework, we expanded our initial implementation of joint cmICA (Wu and Calhoun, 2023) that simultaneously decomposes tractography-based SC (Wu et al., 2015) and temporal-correlation-based FC (Wu et al., 2018, Wu et al., 2021), into a joint decomposition at a finer temporal scale aimed at capturing dynamic alterations in linked patterns between these two different modalities. To achieve this, we ran two separate runs of joint cmICA using the same SC along with FCs assigned with different time frames, segment 1 and segment 2, rather than using the original “static” FC constructed from the entire time series. Subsequently, we employed a similarity measure to match components across runs and identify the degree to which each component of a given modality exhibits static or dynamic characteristics across the two fusions.

**Figure 1.**
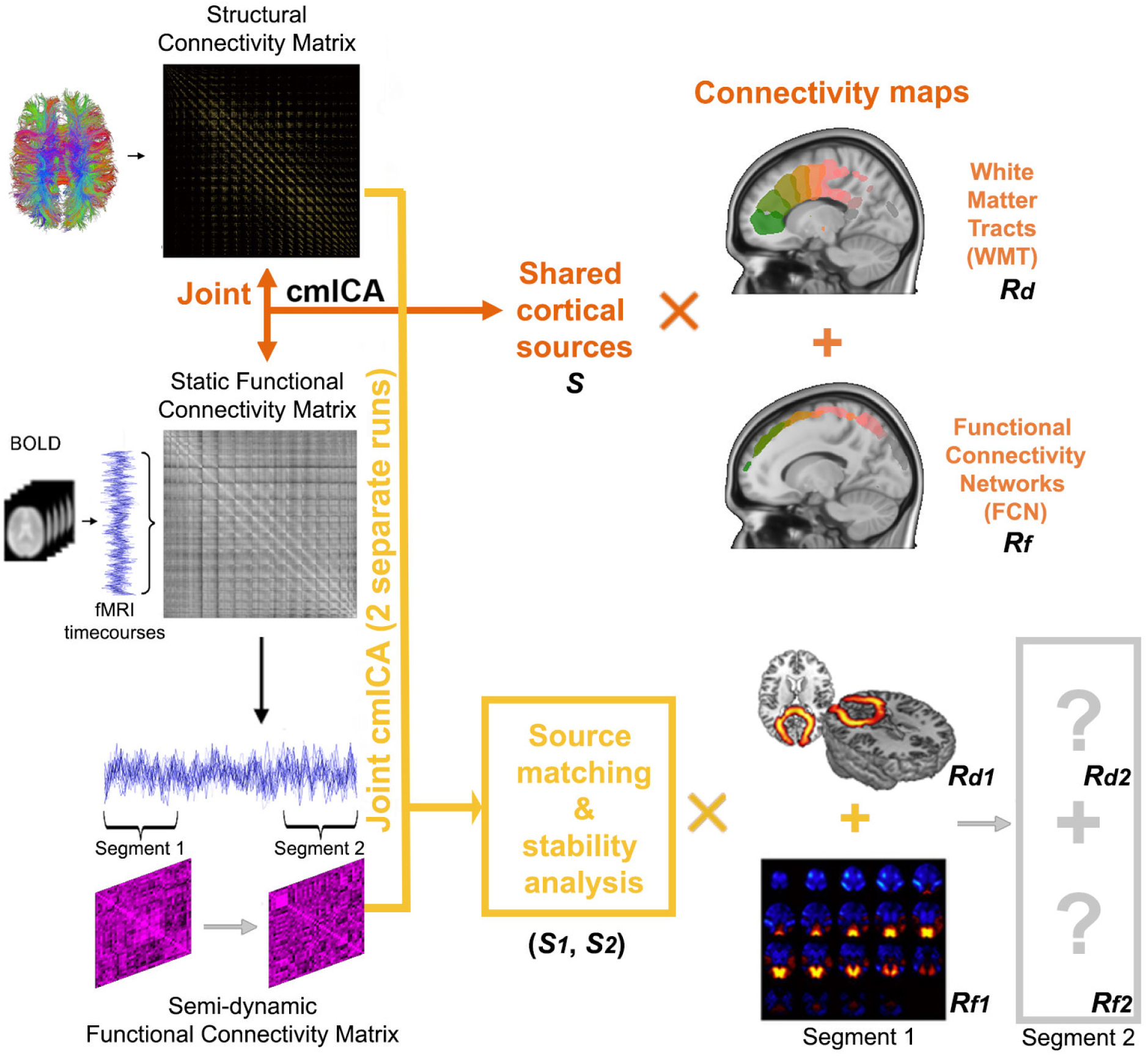
Framework of dynamic fusion. The original joint cmICA separates the SC matrix and FC matrix from the same subject into two parts: shared cortical sources and the connectivity maps corresponding to these sources from both modalities, i.e., how the sources are connected in WMT and FCN, in a single step. The dynamic fusion analysis extends this process to two different time frames and compares the dynamics across two runs of joint separations by analyzing their stabilities.

Mathematically, we can represent the approach as follows. The brain connectivity matrix *C* is parcellated into spatially independent source maps *S* and corresponding connectivity profile maps *R* that define the connectivity of each source *S* towards the whole brain, i.e *C* = *R* × *S* (Wu 2015). Then it is straightforward to derive:

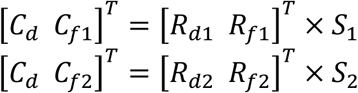

Where *C*_*d*_ is the SC computed from dMRI, and *C*_*f*1_, *C*_*f*2_ are the FC from fMRI with the time window assigned to segments 1 and 2 respectively. *S*_1_, *S*_2_ are the joint sources from two different separations. *R*_*d*1_, *R*_*d*2_, *R*_*f*1_, *R*_*f*2_ are connectivity profile maps from two modalities and two separations. *R*_*d*1,*f*1_ and *R*_*d*2,*f*2_ are pairwise connectivity profile maps from two modalities and two separations. Due to the constraints of the stationarity and independence of ICA, as well as to analyze both white matter tracts (WMT) and functional connectivity networks (FCN) from two different modalities, we primarily focus on comparing the stability of *R* in our framework.

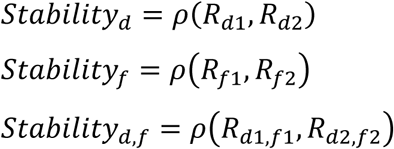

Where ρ represents a sorted similarity function. If we use a normalized distance or similarity, such as absolute correlation, a value of 1 represents a static component, while 0 represents a dynamic/unique component across the two fusions. Note, this framework is generalizable and can be extended to accommodate an arbitrary number of fusions, and it is also possible to perform a joint optimization to solve the problem. However, in this initial work, we illustrate the concept with a two fusion stepwise approach.

Next, we summarize the joint cmICA model used for the fusion process, as well as the similarity measure used to match the two separate joint runs and to quantify the dynamic (i.e., time-resolved) aspect of the fusion.

#### 1.2 Joint cmICA and feature selection

In (Wu and Calhoun, 2023), we combined two cmICAs, one for temporal-correlation-based FC (Wu et al., 2018, Wu et al., 2021, Iraji et al., 2016, Kinsey et al., 2024), and one for tractography-based SC (Wu et al., 2015), into a single algorithm. The joint cmICA performs a blind source separation of gray matter independent sources shared between both modalities (Figure 1). Prior to ICA decomposition, three runs of PCA are conducted, with the first and second runs for dimension reduction at the subject level and group level, respectively, for the FC and SC matrices separately, and the third run applied at the fused level by concatenating both group-level PCAs. This latter data reduction balances the differences in the FC and SC data distributions, avoiding ICA selecting “sources” from one modality only due to the significant differences in variance between FC and SC matrices. Finally, joint cmICA generates connectivity-based cortical sources/parcels which are shared between brain structural and functional connectivity. Their corresponding connectivity maps of functionally connected regions (i.e., functional connectivity networks, FCN) as well as the white matter tracts (WMT) that connected these regions, are jointly identified in one estimation using GICA back-reconstruction and concatenation order.

In this study, we chose a model order of 30 for joint cmICA to effectively capture information from both modalities while maintaining consistency with model dimensions validated in previous work. Specifically, we previously identified 30 reliable cmICA sources in white matter tracts using COBRE dMRI data (Wu et al., 2015). In (Wu and Calhoun, 2023), we employed a bilateral joint cmICA model from two modalities, with a model order of 60 based on the assumption of symmetrical contributions from both fMRI and dMRI connectivity. In the current study, due to the application of an additional fused-level PCA across modalities, we reverted to a model order of 30. To ensure the validity of this choice, ICASSO (Himberg et al., 2004) was used to access convergence and stability during ICA training, selecting the ‘best run’ to ensure robust and replicable results. Joint cmICA yielded 30 independent source maps which were further processed and visualized using the procedures described in (Wu and Calhoun, 2023).

#### 1.3 Similarity measure

In this study, we adapted a multiscale volume-wise similarity measure for 3D images (Wang et al., 2003), for both source matching and stability measure. The applied similarity method computes multiple structural similarity indices between volume A and volume B at various scales and combines them into one score, with a value closer to 1 indicating higher resemblance and a value closer to 0 indicating poorer resemblance. In addition, the similarity method also considers 3D images at different spatial sub-sampling scales. It provides more flexibility than traditional single-scale methods (e.g. spatial correlation) in incorporating the variations of viewing conditions and is closer to human visual sensitivity (Wang et al., 2003).

Using this measure, we evaluated whether the sources generated from different separations were well-matched. Once the sources were determined to be matched, we employed this measure to assess the stability/dynamics along connectivity maps R using the same matching pair indices.

Additionally, we explored various applications of such methods in our study. Specifically, we employed 1) a single overall similarity score for component matching and sorting, averaging across volume and scale-weighted local indices for each sampling volume, and 2) 3D volumes of similarities and dissimilarities (i.e., 1-*ρ*) at voxel scale to identify subtle spatial differences across states and subject groups.

Next, we show an application of the proposed approach to a real data set including fMRI and dMRI data collected from 117 individuals.

### 2. Subjects

Subjects were recruited via a Center Of Biomedical Research Excellence (COBRE, http://cobre.mrn.org) program at the Mind Research Network (MRN). In this paper, we used fMRI and dMRI data from a large set of schizophrenia (n = 57, age = 39.0 ± 13.4 years) and typical control subjects (n = 60, age = 36.8 ± 12.1 years), which we previously studied to estimate SC and FC separately in unimodal cmICA analyses (Wu et al., 2015, Wu et al., 2018, Wu and Calhoun, 2023). In this case, we excluded two individuals with schizophrenia and one control subject who did not have both imaging modalities collected. All the participants were enrolled in the COINS platform (http://coins.trendscenter.org) (Scott et al., 2011, Wood et al., 2015). Prior to inclusion in the study, subjects were screened to ensure they were free from neurological or psychiatric disorders (DSM-IV Axis I) as well as active substance use disorders (except for nicotine).

### 3. Data acquisition and preprocessing

Data were collected and preprocessed as described in (Wu et al., 2018, Wu et al., 2015), with additional graphics processing unit (GPU) computational upgrades. All participants were scanned on a 3Tesla Siemens TIM Trio equipped with a 12-channel radio frequency coil. Subjects were instructed to fixate on a central crosshair presented with eyes open at rest.

Diffusion data were acquired via a 6-minute single-shot spin-echo echo planar imaging (EPI) with a twice-refocused balanced echo sequence (30 directions, b=800 s/mm^2^,5 b0, Field of view (FOV) = 256 × 256 mm, slice thickness = 2 mm, slices = 72, matrix size = 128x128, voxel size = 2 × 2 × 2 mm^3^, echo time (TE) = 84 ms, repeat time (TR) = 9000 ms, number of excitations (NEX) = 1, partial Fourier encoding = 3/4, GRAPPA acceleration factor = 2). Preprocessing was performed using FSL (http://fsl.fmrib.ox.ac.uk), including motion and eddy current corrections, with directions showing excessive signal dropout excluded from analysis. ODF diffusion parameters were estimated using bedpostx GPU, and whole-brain probabilistic tractography (2000 streamlines/voxel) was conducted via probtrackx GPU without predefined ROIs. Data were downsampled to 5 mm resolution (14,811 nodes) to manage matrix size. Final voxelwise connectivity profiles were compiled into a 2D matrix representing fiber counts between voxel pairs.

Resting state functional scans were collected using 5-minute T2*-weighted gradient echo planar imaging (TR = 2 s, TE = 29 ms, FOV = 240 mm, acquisition matrix = 64×64, flip angle = 75°, voxel size = 3.75 × 3.75 × 4.55 mm^3^, gap = 1.05 mm, 33 slices, ascending acquisition). FMRI data preprocessing with SPM included discarding the first five volumes for T1 equilibration, realignment using INRIalign, slice timing correctio, normalization to MNI space, smoothing (FWHM = 5mm), and sub-sampling to 5x5x5 mm³ to match dMRI data resolution. Further steps involved detrending, despiking, and band-pass filtering (0.001-0.15 Hz) to remove noise (Wu et al., 2018, Allen et al., 2011, Cordes et al., 2000). White matter and gray matter masks were used for facilitating accurate cortical parcellation and white matter tract extraction (Wu and Calhoun, 2023). Finally, temporal normalization was applied to each voxel to optimize cmICA computation (Wu and Calhoun, 2023).

Granted, any time segments could be used in our dynamic fusion framework (Figure 1), but to enhance temporal separation between the two fusions, we explored the first and the last third of the entire time series as exemplary segments in the current study, and these segments are referred to as “States”.

## Results

### 1. Joint cortical parcellations at two different time frames

As described in the methods section (Figure 1), we utilized shared-sources *S* from two separate joint cmICA results for component matching, leveraging their within-modality spatial distribution similarity in comparison to the *R* maps (Wu, 2018, 2023).

Figure 2 illustrates the matched shared-sources *S*, sorted from highest to lowest similarity. In this figure, State 1 is represented in red, while State 3 is shown in blue; the areas of overlap are depicted in a purple hue. Notably, most components demonstrate a strong degree of matching. Among the 30 joint source matches, only two pairs exhibited 3D similarity scores below 0.6 (specifically, ρ = 0.56 and 0.44). This indicates that the majority of the components are highly consistent, supporting our hypothesis that cortical sources exhibit a significant degree of stability overall. However, there is also a notable level of dynamics and flexibility observed across the two different states. This evidence reinforces the validity of our framework for analyzing dynamic changes in connectivity.

**Figure 2.**
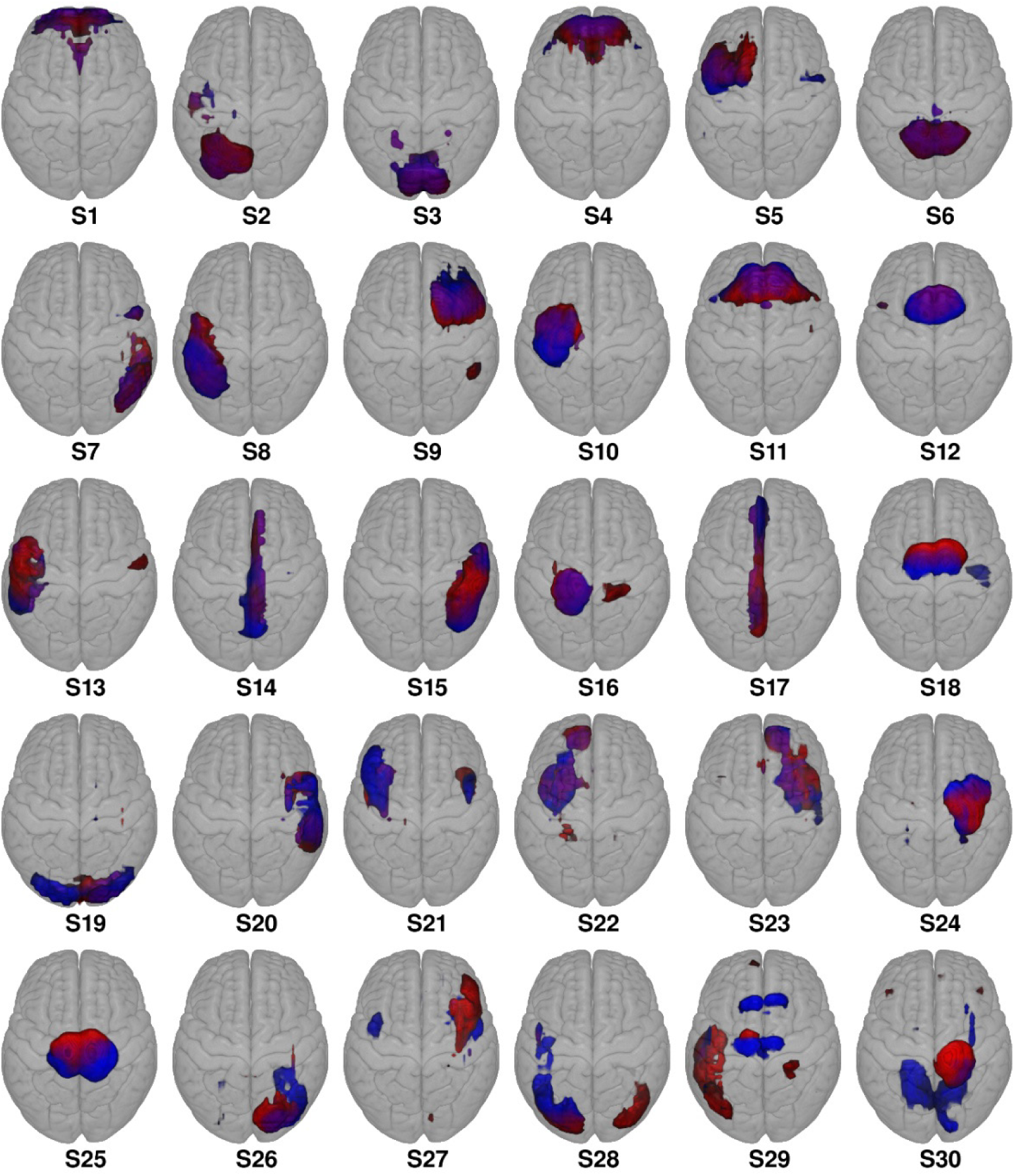
Matching joint cortical sources. The figure shows two runs of shared sources (S_1_ and S_2_ respectively, matched and sorted by 3D similarity. Most components exhibit high similarity, with only two pairs scoring below 0.6 (S29 = 0.56 and S30 =0.44).

### 2. Sorting

All other component WMTs and FCNs (*R_d_*, *R_f_*) are matched following the index order established in the joint cortical sources *S* mentioned above. Using these matches, we compared the similarity rankings for all relevant components, including shared *S* maps, *R* maps for each modality, and a dual similarity across the two modalities. The results found that the similarities measured across these components generally, but not perfectly, follow a similar high-to-low direction of stability as observed in the shared *S* maps.

To clarify these relationships, we sorted the similarities in their respective orders. Additionally, due to the differing order between the structural *R_d_* and functional *R_f_* maps, we computed a dual similarity by averaging the similarities from pairwise *R_d_* and *R_f_* in joint cmICA. This allowed us to sort the dual similarities, enabling us to observe dynamic changes within the same joint component more effectively.

Figure 3 displays the within modality similarity (from high to low) sorted separately. The solid green line represents the stability of the shared *S* maps across State 1 and State 3. The blue dashed line indicates the stability of the SC (WMT) *R_d_* maps between the two States. The red dashed line reflects the stability of the FC (FCN) *R_f_* maps between the two States. Finally, the violet dashed line shows the stability of the dual similarities derived from the pairwise *R_d_* and *R_f_* maps from joint cmICA across the two States. Below, we use the sorting order based on the dual similarity.

**Figure 3.**
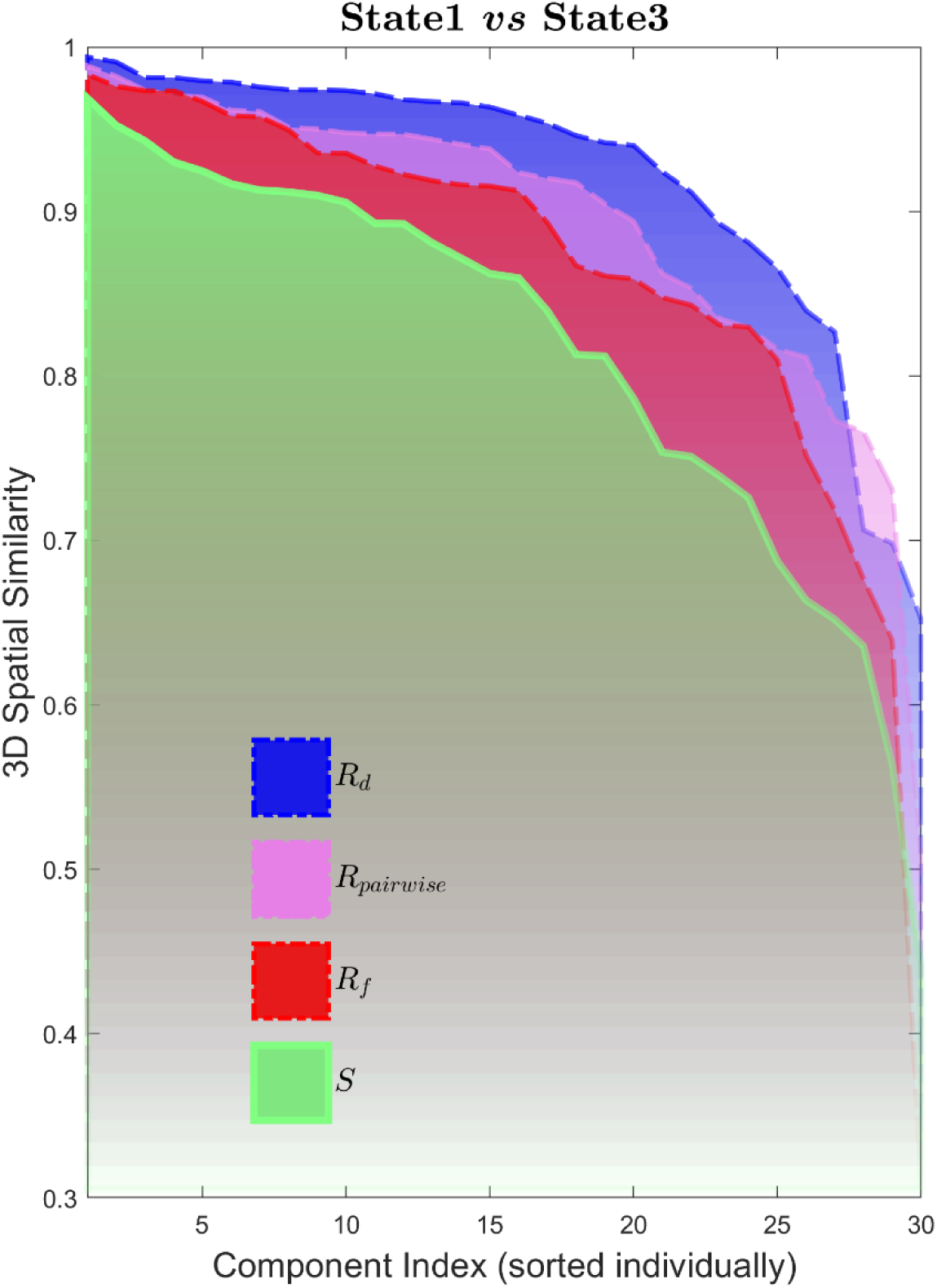
Cross-Fusion Similarity. This figure displays within- and cross-modality similarities, sorted in descending order respectively. The solid green line shows the stability of shared *S* maps across State 1 and State 3, while the blue and red dashed lines represent SC *R_d_* and FC *R_f_* maps. The violet dashed line indicates pairwise dual similarities from both *R_d_* and *R_f_*.

### 3. Stable and dynamic multimodal components

In this section, we explore the distinction between stable and dynamic *R* maps within the framework of our analysis, by focusing on the five components with the highest and lowest similarity. Figure 4 and 5 show the top five stable and dynamic multimodal components, respectively, with yellow indicating State 1 and blue indicating State 3, and cyan hue indicating the overlaps between the two States.

**Figure 4.**
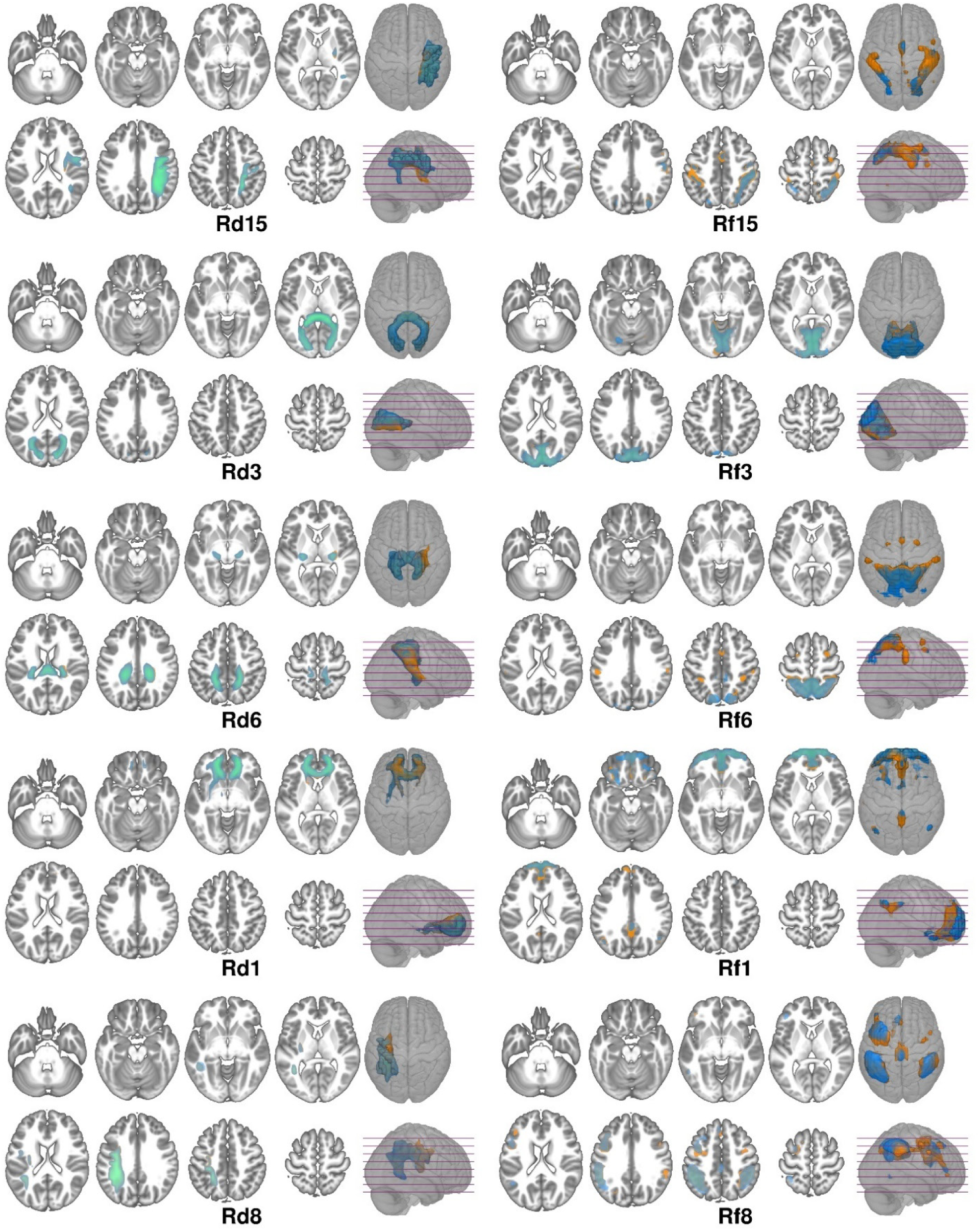
Stable multimodal components. The plots display the top five components with the highest 3d similarity scores across States, indicating consistent connectivity patterns, with yellow indicating State 1, blue indicating State 3 and cyan hue indicating the overlaps. These components are vital for understanding brain relationships as they reveal essential connectivity patterns that underpin baseline brain functions that are more resilient over time and in mental processes.

**Figure 5.**
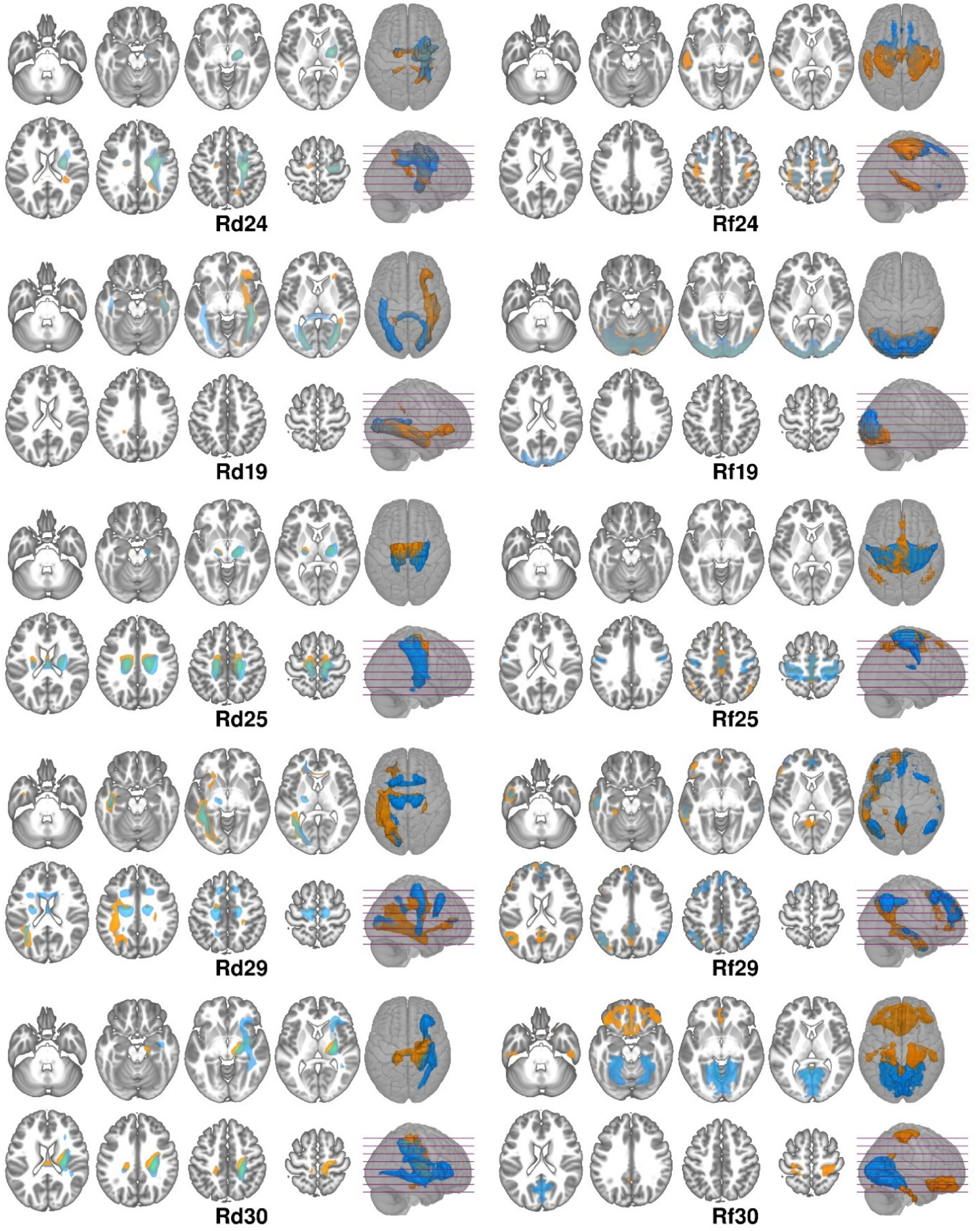
Dynamic multimodal components. The plots display the top five components with the lowest 3d similarity scores across states, indicating dynamic connectivity patterns, with yellow indicating State 1, blue indicating State 3 and cyan hue indicating the overlaps. These components are crucial for identifying and understanding how different regions interact and transition over time and across different mental functions (e.g. drowsiness, fatigue).

Stable multimodal components are characterized by high similarity scores across the two States, indicating a consistent pattern of connectivity that persists over time. These components are crucial for understanding the foundational structural and functional relationships in the brain, highlighting the connectivity patterns that support baseline brain functions where it exhibits greater stationarity in temporal and mental contexts. Figure 4 presents the top five components with the highest 3D similarity scores across States. The results indicate stability in the commissural tract bundles that radiates from two ends of the corpus callosum (the splenium, *R_d3_*, and the genu *R_d6_*, *R_d1_*), which connect the occipital (*R_f3_*), parietal (*R_f6_*), and prefrontal (*R_f1_*) regions. Notably, *R_f1_* reveals a slight transition in the DMN, with State 1 (yellow) showing a stronger connection between the prefrontal region and the PCC compared to State 3 (blue). Additionally, the right (*R_d15_*) and left (*R_d8_*) superior longitudinal fasciculi, linking to the dorsal attention network (*R_f15_*) and motor network (*R_f8_*), are relatively stable as well.

Conversely, dynamic multimodal components exhibit greater variability in their similarity scores, possibly reflecting the brain’s capacity to adapt and respond to different brain conditions, status and/or vigilance. By identifying these dynamic multimodal components, we can gain insights into how neural connectivity shifts in response to varying cognitive demands or environmental changes. Figure 5 illustrates significant dynamic changes in both WMT and FCN patterns, particularly in *R_d,f19_*, *R_d,f29_* and *R_d,f30_*. However, these dynamics do not necessarily behave the same and exhibit an even distribution between WMT and FCN. For example, the FCN *R_f19_* shows strong local connectivity to source *S_19_*, located near the visual and occipital areas in both States. In contrast, the WMT *R_d19_* demonstrates a shift in connectivity, transitioning from connections through the forceps major across the corpus callosum, the inferior fronto-occipital and inferior longitudinal fasciculi to a predominant focus on the forceps major and inferior longitudinal fasciculus.

Note that connectivity maps, *R_f_* particularly, in Figures 4 and 5 do not always perfectly overlap with their corresponding sources *S* in Figure 2, as noted in (Wu et al., 2018). *R_f_* reflects whole-brain FC of each source *S*, highlighting both intra- and inter-regional patterns. In general, *R* maps are more diffuse compared to their matched *S* maps. High overlap (e.g., *S_3_*/*R_f3_*, *S_6_*/*R_f6_*) indicates strong intra-source connectivity, while lower overlap (e.g., *S_29_*/*R_f29_*, *S_30_*/*R_f30_*) suggests broader inter-regional links. Consistent with our earlier findings, *R* maps also reveal strong bilateral and inter-DMN connectivity, in contrast to the original *S* maps which are split into unilateral or partial-DMN components, such as S_15_/*R_f15_*, *S_8_*/*R_f8_*, *S_24_*/*R_f24_*, *S_1_*/*R_f1_* and *S_29_*/*R_f29_*.

Our analysis reveals that while many components remain stable, a subset of the estimated dMRI components display significant dynamism (i.e., time-dependent fusion). This variability may be important for determining both context-independent and context-dependent structural substrates of state-based brain networks, for example where cognitive flexibility is required, such as during shifts in vigilance or responses to different brain status or stimuli. By examining both stable and dynamic multimodal components, we can better understand the underlying mechanisms of brain function and connectivity, ultimately contributing to a more comprehensive view of neural dynamics.

### 4. Group differences

To compare group differences between controls (HC) and schizophrenia (SZ), and to maintain consistency with our adopted similarity process, we developed a two-step method. First, we examined the similarity between States on *R* components using the same sorting index derived from dual similarities, computed separately for HC and SZ. Second, we retained the initial 3D similarity volume maps across State 1 and State 3 for both HC and SZ, denoted as Ρ_hc_ and Ρ_sz_ (as described in Methods 1.3 Similarity measure), rather than a single overall similarity score from the first step, and conducted another round of 3D similarity assessments now between HC and SZ using their similarity voxel maps across States, i.e., *Stability*_*hc,sz*_ = *ρ*(*P_hc_, P_sz_*).

Figure 6 presents the five most stable and the five most dynamic multimodal components. The blue bars illustrate the 3D similarity between State 1 and State 3 for HC, while the gray bars represent the similarity for SZ. Purple bars indicate the 3D similarity between the HC and SZ groups, as defined above. The left subplot focuses on SC similarity, highlighting how stable and dynamic components interact within the brain’s structural framework. In contrast, the right subplot addresses FC similarity, providing insights into the static and dynamic interactions that occur during different times and/or mental states. Figure 6 shows a clear drop in stability from left to right for HC and SZ individually, as well as for similarity between HC and SZ, in both SC and FC. The decline in FC is more pronounced due to its higher variability. Interestingly, HC exhibits less decrease of stability in FC compared to SZ, which we speculate may suggest that HC maintain vigilance more effectively, however future studies would have to test this directly.

**Figure 6.**
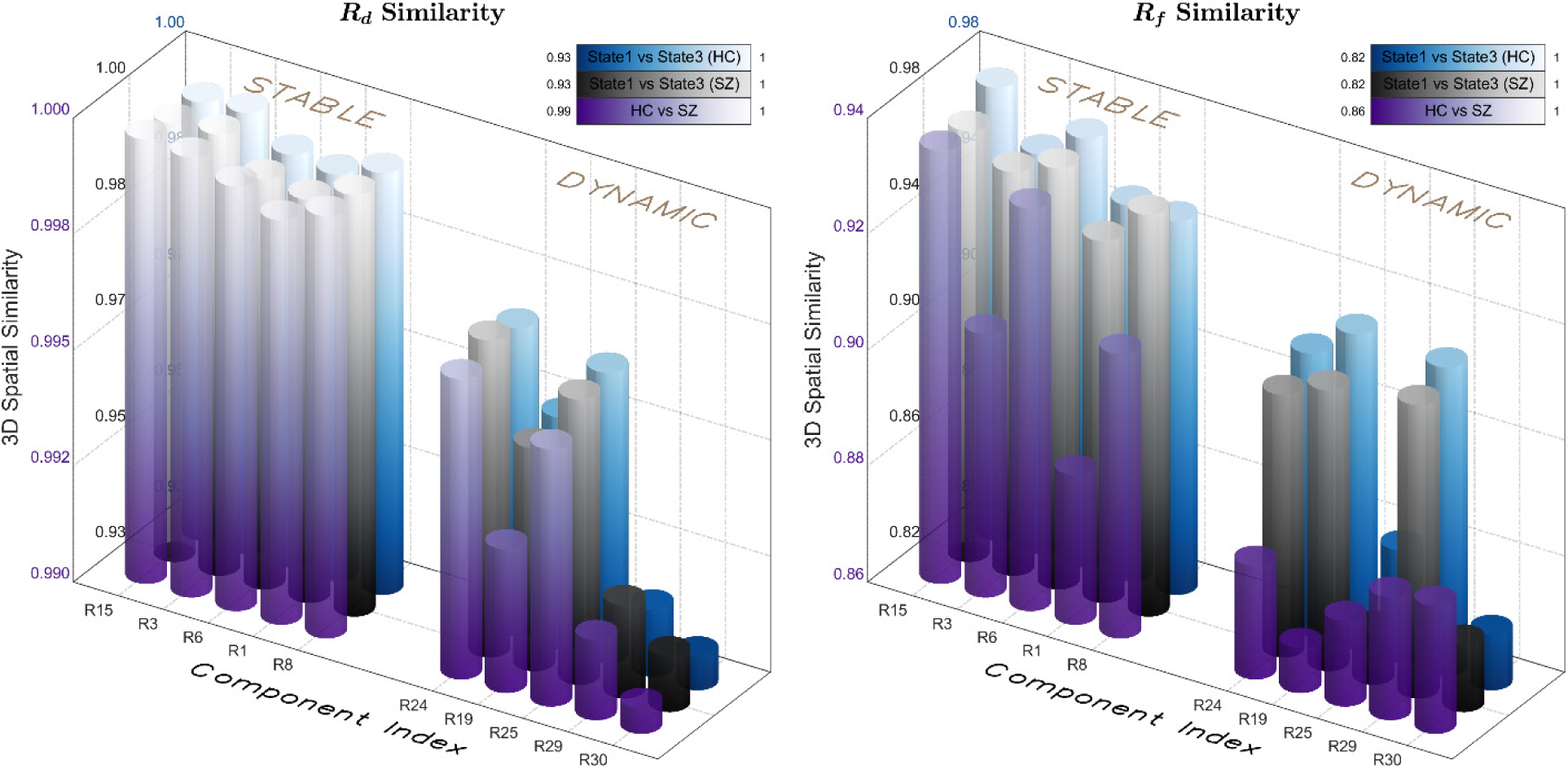
Top 5 most stable and most dynamic multimodal components. Blue bars represent 3D similarity between State 1 and State 3 in HC, gray bars indicate the similarity in SZ, and purple bars show 3D similarity between HC and SZ. The left subplot focuses on SC, while the right subplot addresses FC similarity. As expected, structural components are more stable overall, but interesting they also can be subdivided into highly stable (i.e., context-independent fusion) and more dynamic (i.e., context-dependent fusion) components.

This comparative approach allows for a clearer understanding of how stability and dynamism manifest in both control and schizophrenia clinical populations, shedding light on potential differences in brain connectivity patterns. By examining these components, researchers can gain valuable insights into the underlying mechanisms of brain function and the impact of conditions such as schizophrenia.

To enhance clarity in presenting these differences, we chose to display the dissimilarity (1 - ρ) for examined, only two dynamic components showed significant group dissimilarity between HC and SZ in both *R_d_* and *R_f_* maps (cluster criteria: Rd: dissimilarity > 0.35, size > 250mm³; Rf: dissimilarity > 0.65, size > 250mm³; smoothed to 6mm). The remaining components showed group differences either in *R_f_* only or in neither modality. Figure 7 illustrates the group dissimilarities for these two dynamic components. See Figure 8 and Supplemental for more details on all ten components.

**Figure 7.**
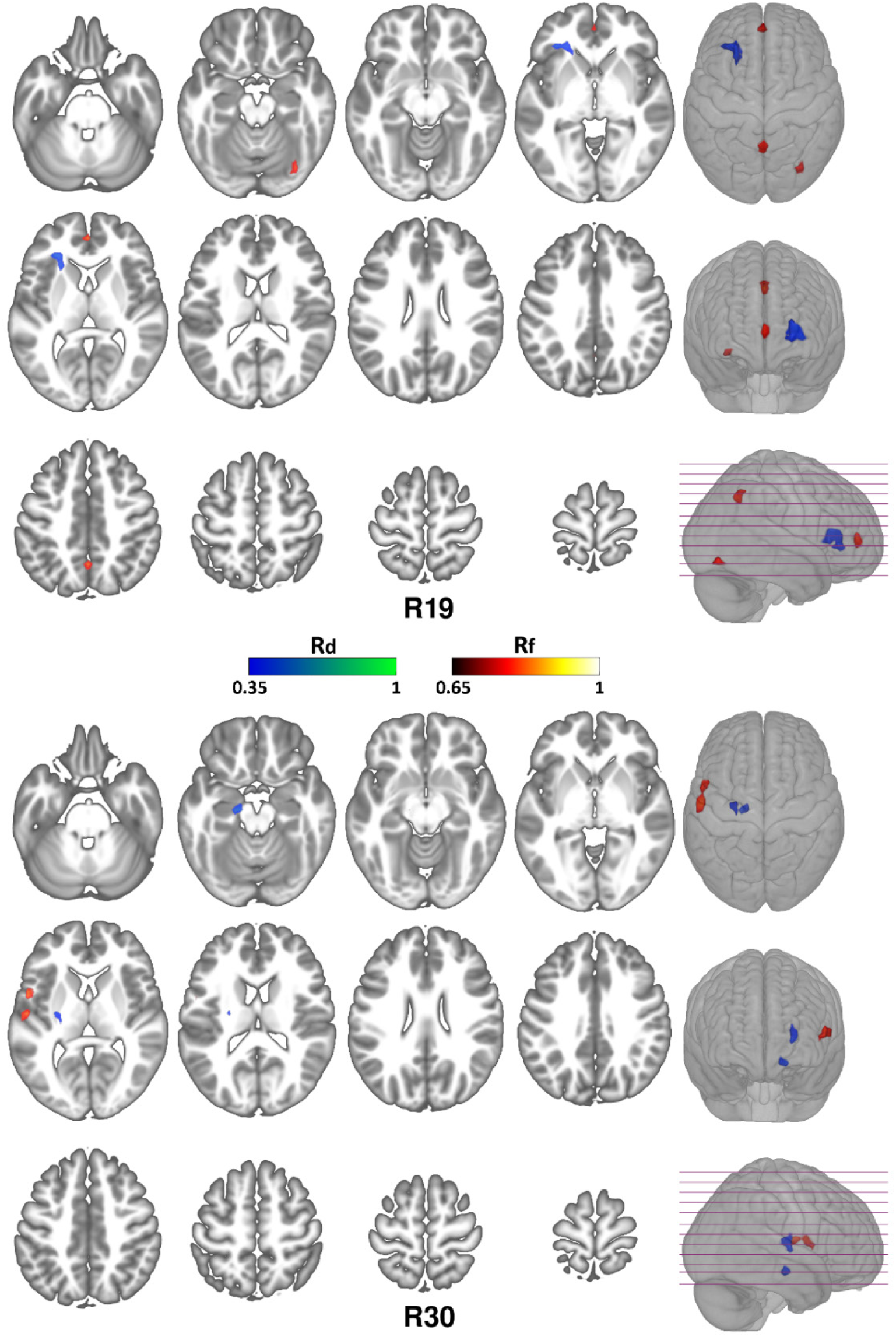
Group dissimilarity of dynamic multimodal components. This figure highlights a much greater dissimilarity in SC between HC and SZ across States, compared to stable components (See Supplemental), with blue to green indicating *R_d_* and red to yellow indicating *R_f_*. Notably, *R_d19_* and *R_d30_* exhibit significant structural dissimilarities, particularly in the anterior thalamic radiation/inferior fronto-occipital fasciculus, and corticospinal tract/anterior thalamic radiation, respectively. Additionally, substantial dissimilarities are observed in the anterior cingulate/paracingulate/precuneus/occipital lobe and precentral/inferior frontal/parietal gyrus, as indicated by *R_f19_*, *R_f30_*.

**Figure 8.**
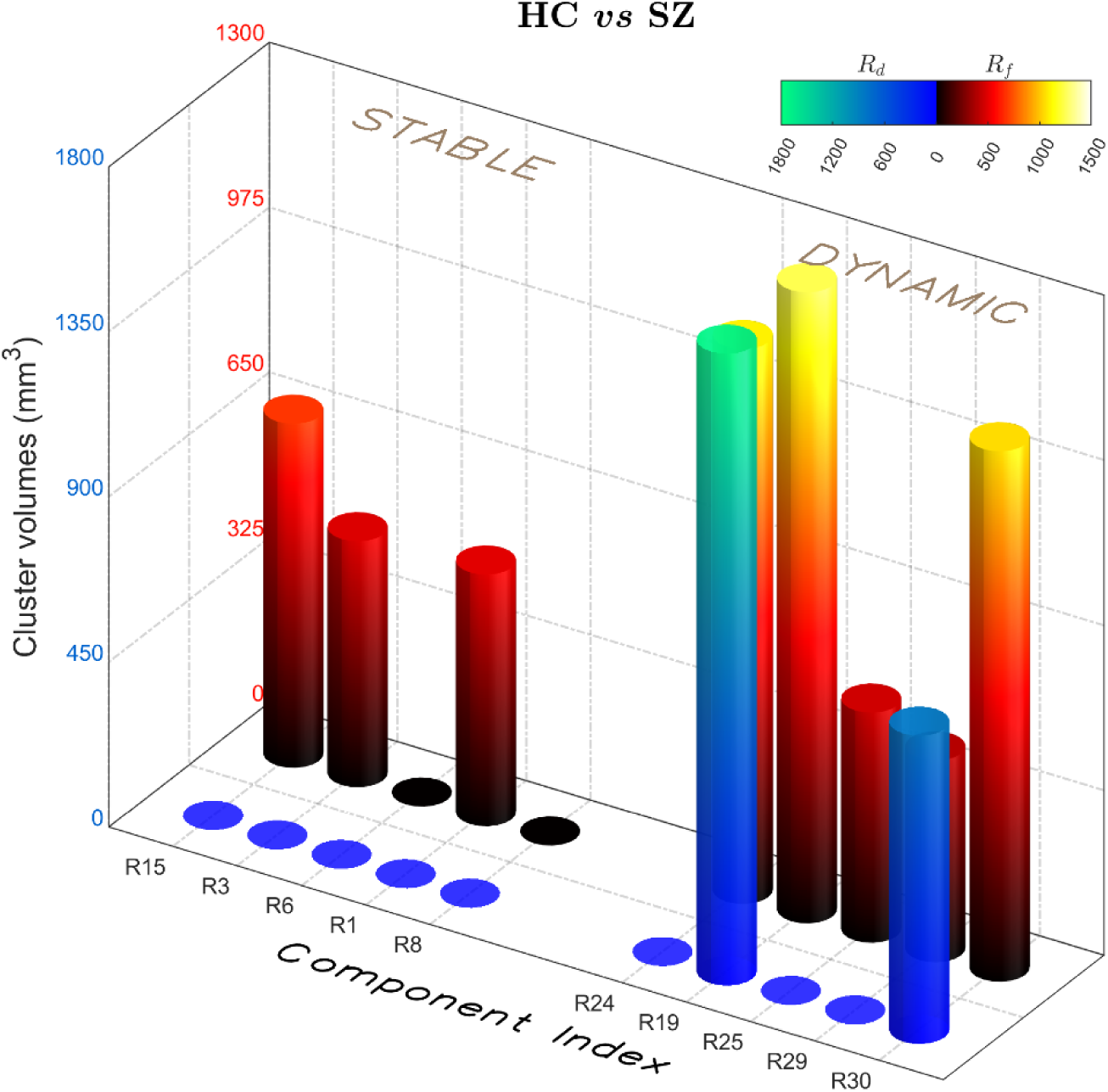
Bar plot of component clusters. It shows the overall volumes of cluster in each component for the top 5 stable and dynamic multimodal components, with blue to green indicating *R_d_* and red to yellow indicating *R_f_*. The details of each clusters are listed in Table 1 in Supplemental.

Overall, the findings in Figure 7 and 8 reveal significantly less variation between HC and SZ in the stable components compared to the dynamic components for both SC and FC. This suggests that the stable multimodal connectivity patterns remain not only relatively consistent across temporal states but also across participant groups, whereas the dynamic multimodal connectivity patterns show both greater temporal variability and pronounced differences between HC and SZ. These differences may reflect underlying disruptions in the fluctuating neural networks associated with SZ, indicating a potential impairment in the adaptive responses to changing cognitive demands.

## Discussion

### 1. Temporal dynamics of joint connectivity networks

While prior studies have focused on linking SC to resting fMRI (Calhoun and Sui, 2016), these approaches typically do not compare across fusions to evaluate the degree to which the multimodal components extracted may vary as a function of the fMRI dynamics. More specifically, to our knowledge, there have been no studies of the degree to which this temporal variation is associated with SC components in a joint decomposition. This study provides an opportunity to evaluate whether different structural sources vary (or do not vary) as a function of the data to which it is fused. The results of this study provide compelling insights into the dynamics of cortical connectivity across different brain states, particularly highlighting the distinctions between HC and SZ. Our findings reinforce the notion that while certain aspects of brain connectivity exhibit remarkable stability, significant dynamism exists, reflecting the brain’s adaptability to varying cognitive demands and environmental.

### 2. Stable multimodal connectivity

The strong correspondence between joint cortical shared sources *S* (Figure 2) and the extensive matching sets of joint connectivity maps *R* (Figure 4) across different states reveals a fundamental stability in both cortical functional connectivity and the underlying white matter tract architecture. Most source components displayed high similarity scores, with only two pairs falling below the 0.6 threshold. This consistency suggests that certain structural and functional connections persist across different mental states, potentially supporting baseline brain functions. The stability of components, particularly those associated with key networks such as the default mode network (DMN) and dorsal attention network, highlights the importance of these pathways in maintaining cognitive and perceptual processes. Notably, the stable multimodal components identified in Figures 4 indicate that critical commissural tracts, such as those radiating from the corpus callosum, remain robust across varying conditions. These connections are vital for interhemispheric communication and play a crucial role in integrating sensory information and coordinating higher-order cognitive functions.

### 3. Dynamic multimodal connectivity

The analysis of dynamic multimodal components (those that showed context-dependent variation across fusions) revealed significant variability in connectivity patterns, within both the SC (WMT) and FC (FCN) framework. Figures 5 maps and Figure 6 (blue and gray bars in the right ‘DYNAMIC’ section) illustrate that dynamic multimodal components exhibit greater fluctuations in similarity scores, which may reflect the brain’s ability to adapt to changing cognitive demands or states of alertness. This variability is crucial for understanding the neural mechanisms underlying cognitive flexibility, as it allows for rapid reconfiguration of networks in response to different tasks or environmental stimuli. The pronounced differences in dynamic connectivity between HC and SZ groups suggest that the latter may experience disruptions in their adaptive neural responses. For instance, the marked dissimilarity observed in Figures 6 and 7, particularly in SC, indicates that SZ subjects may struggle with maintaining stable and adaptive connections, which could contribute to the cognitive impairments and symptoms associated with the disorder. The inability to modulate connectivity patterns effectively may impair the brain’s ability to respond flexibly to external challenges, potentially exacerbating symptoms such as attention deficits and executive function difficulties.

### 4. Dynamic connectivity fusion in schizophrenia

Our dynamic connectivity fusion model provides a more comprehensive framework for distinguishing between healthy and disordered brain states. By integrating both stability and variability across structural and functional domains, the model captures connectivity disruptions more effectively. SZ individuals exhibit greater instability and variability in both Rd and Rf the development of targeted, personalized interventions and is especially useful for identifying patterns in disorders that are not well characterized by traditional, stationary analyses.

This study offers valuable insights into the neural underpinning of schizophrenia. While some SC and FC patterns appear stable, the increased variability in dynamic components suggests reduced network flexibility in individuals with SZ. Since this flexibility is critical for adaptive cognitive functioning, its disruption may contribute to the cognitive deficits commonly observed in this population. These findings underscore the importance of assessing both stable and dynamic multimodal connectivity to fully understand brain function in clinical populations.

### 5. Limitation

#### 5.1 Two Separate Runs of Joint cmICA

One significant limitation of the current study is the reliance on two separate runs of joint component mutual information analysis (cmICA) instead of employing a unified dynamic cmICA joint model. While this methodological choice allows for a focused examination of distinct brain states, it inherently limits the continuity and fluidity of the connectivity patterns being analyzed (Iraji et al., 2020, Iraji et al., 2019). Each run captures a snapshot of the brain’s FC at a specific time frame, potentially neglecting the dynamic nature of neural interactions that occur in real time. Additionally, State-wise differences at subject level require further examination.

This segmentation can lead to an over-reliance on the results derived from “dynamic” segment assignments. The accuracy of the current joint links and difference detection becomes contingent upon the fidelity of these segments. If the temporal segment/assignment calculations fail to accurately represent the underlying neural dynamics, this could compromise the validity of the findings. Additionally, the lack of integration across runs may hinder the identification of transient or subtle connectivity changes, which are crucial for understanding the complexity of cognitive processes and the pathophysiology of conditions such as schizophrenia.

Future studies should consider integrating the findings from multiple runs into a more cohesive dynamic cmICA framework. This approach could enhance the ability to track continuous changes in connectivity and provide a richer understanding of how brain networks interact over time.

#### 5.2 Dissimilarity measurement

In this study, we used 1 - ρ as a dissimilarity measure to compute the differences between groups. However, it is important to recognize that the adapted 3D multiscale similarity measure we employed does not conform to essential mathematical properties required for a valid distance function, namely the triangle inequality and non-negativity (Brunet et al., 2011). As a result, the values generated through this approach do not genuinely represent a “distance” between components, complicating our interpretation of group differences and potentially raising concerns regarding the robustness of our dissimilarity measure.

In addition, the negative values in our dissimilarity measure poses additional interpretive challenges. Negative dissimilarity values can obscure the distinction between groups, making it difficult to identify and articulate the specific nature of the differences observed. Employing metrics that satisfy the triangle inequality and non-negativity would provide a clearer foundation for interpreting group differences and enhance the reliability of our conclusions. Additionally, validating the chosen dissimilarity measure against established standards in the field could yield insights into its effectiveness and limitations, ensuring that subsequent analyses are both rigorous and meaningful. By addressing these methodological concerns, we can improve the interpretability of our findings and contribute to a more nuanced understanding of the neural dynamics underlying various cognitive and clinical states.

## Conclusion

In summary, this study introduces a novel way to analyze dynamics between structural and functional connectivity and highlights the dual nature of cortical connectivity—combining both stability and dynamism—as essential for understanding brain function. The distinct patterns observed between HC and SZ populations emphasize the need for further research into the adaptive mechanisms of neural networks and their implications for cognitive health. Future studies should continue to explore the interplay between stable and dynamic connectivity, particularly in clinical contexts, to identify potential targets for intervention and enhance our understanding of psychiatric disorders.

