## Supplemental Table 1 and Figure S1 to S10 for "Dynamic fusion of structural and functional connectivity via joint connectivity matrix ICA"

1.Table 1. Dissimilarities between HC and SZ across States

| Structural | Cluster volume (mm$^3$) | Peak XYZ (mm) | Functional | Cluster volume (mm$^3$) | Peak XYZ (mm) |
|---|---|---|---|---|---|
| | | ***Stable multimodal components*** | | | |
| Rd15 | 0 | N/A | Rf15 | 311 | 5.4×-82.7×-13.5 |
| | | | | 366 | 29.0×-79.1×-17.2 |
| Rd3 | 0 | N/A | Rf3 | 484 | 48.9×-35.5×49.2 |
| Rd6 | 0 | N/A | Rf6 | 0 | N/A |
| Rd1 | 0 | N/A | Rf1 | 495 | -52.9×2.8×0.5 |
| Rd8 | 0 | N/A | Rf8 | 0 | N/A |
| | | ***Dynamic multimodal components*** | | | |
| Rd24 | 0 | N/A | Rf24 | 1092 | -4.2×-97.5×13.8 |
| Rd19 | 1722 | -27.1×30.8×4.2 | Rf19 | 539 | -0.5×-57.7×44.8 |
| | | | | 423 | -1.3×50.0×4.2 |
| | | | | 280 | 33.4×-76.1×-15.0 |
| Rd25 | 0 | N/A | Rf25 | 453 | -53.6×2.8×-12.0 |
| Rd29 | 0 | N/A | Rf29 | 397 | 1.7×-1.6×49.9 |
| Rd30 | 308 | -17.5×-14.9×-16.5 | Rf30 | 1044 | -55.9×-4.6×10.8 |
| | 532 | -26.4×-12.7×9.4 | | | |

* Cluster size: $R_d$ > 250mm$^3$, $R_f$>250mm3, smoothed to 6mm.

2. Stable and dynamic multimodal components

Stable multimodal components (Figure S1-S5). It shows no difference in structural connectivity between HC and SZ across two States for the top five stable components, but slight changes in functional connectivity, particularly in $R_{f15}$, $R_{f3}$ and $R_{f1}$.

Figure S1.

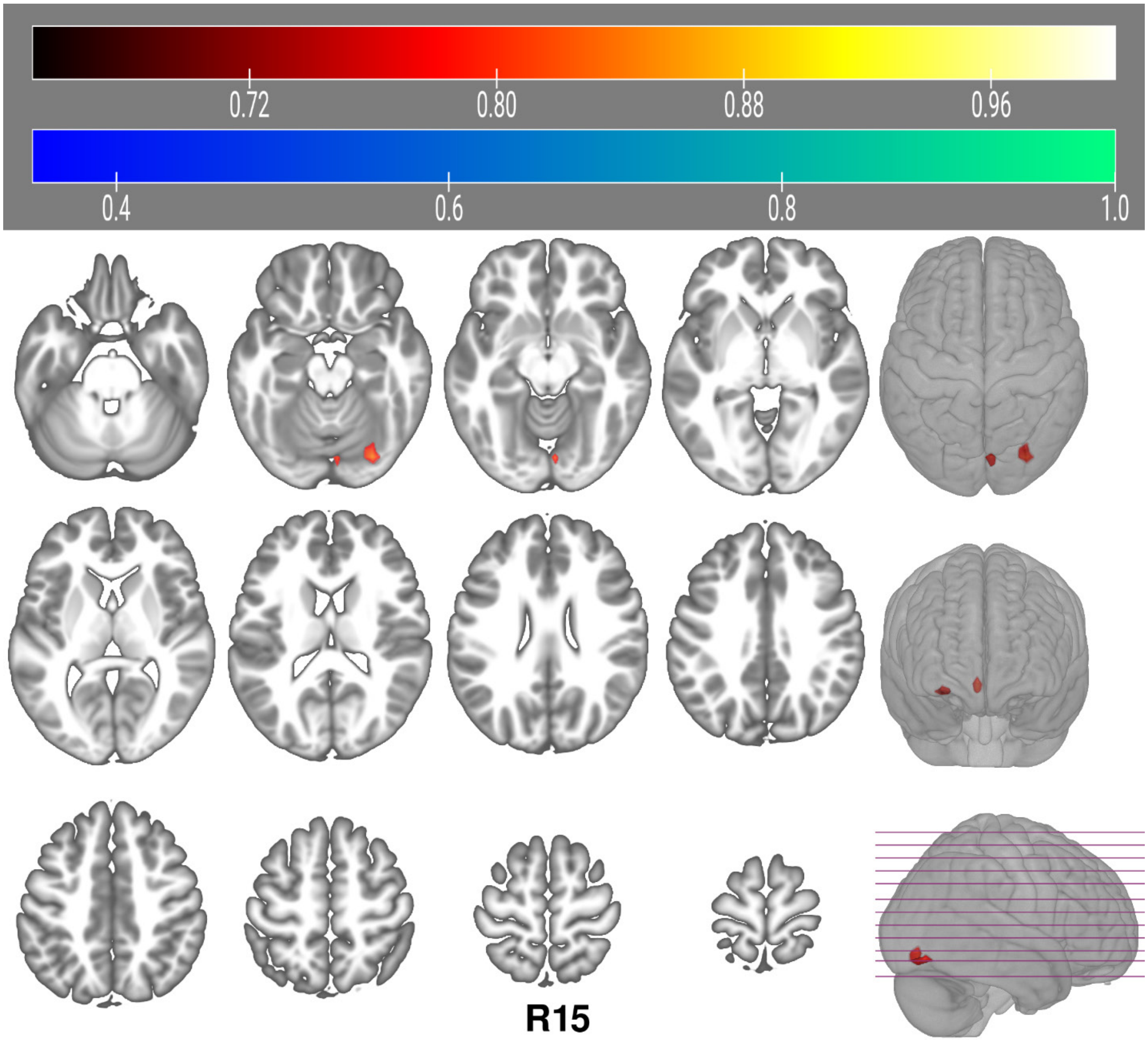

Figure S2.

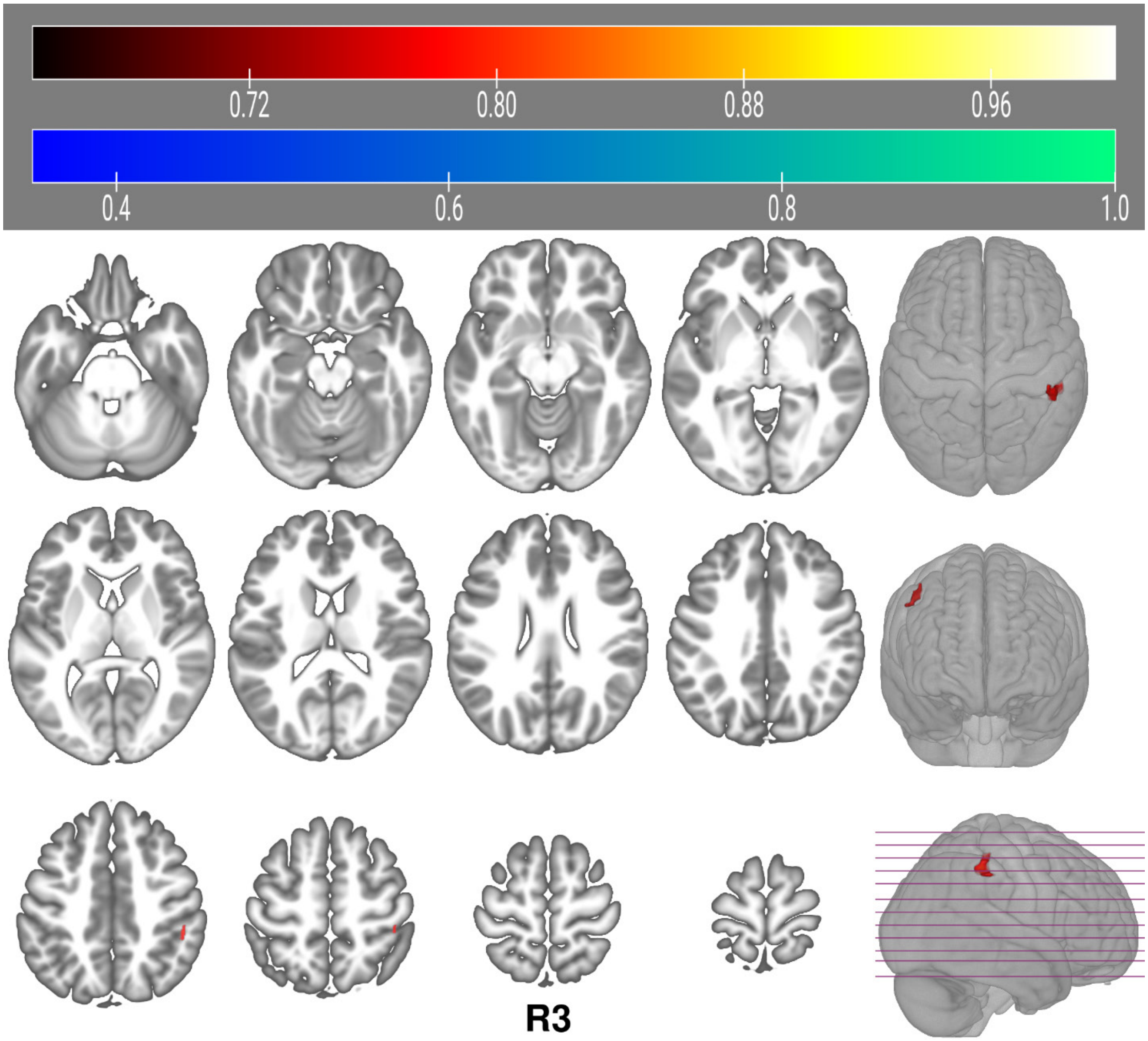

Figure S3.

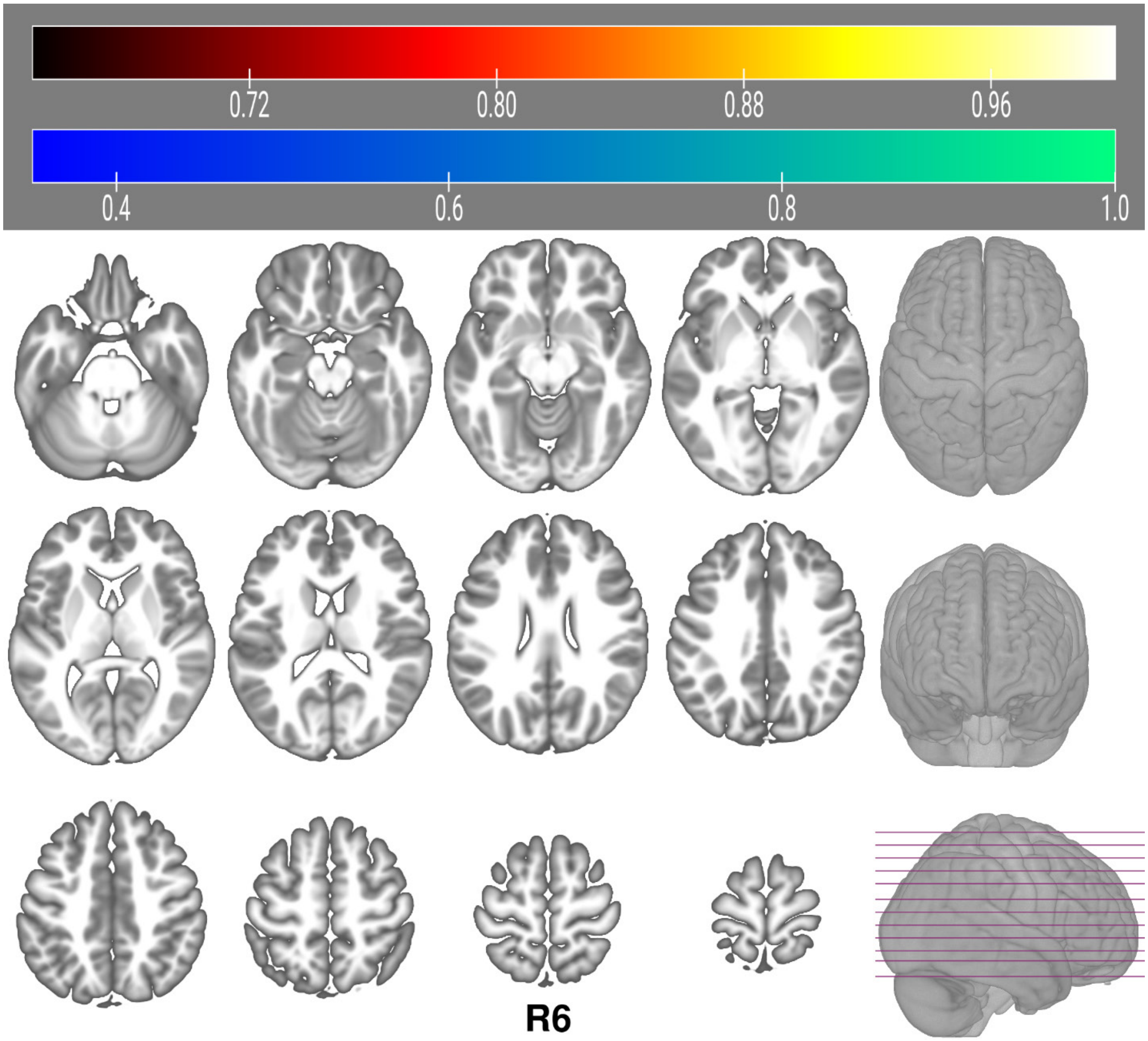

Figure S4.

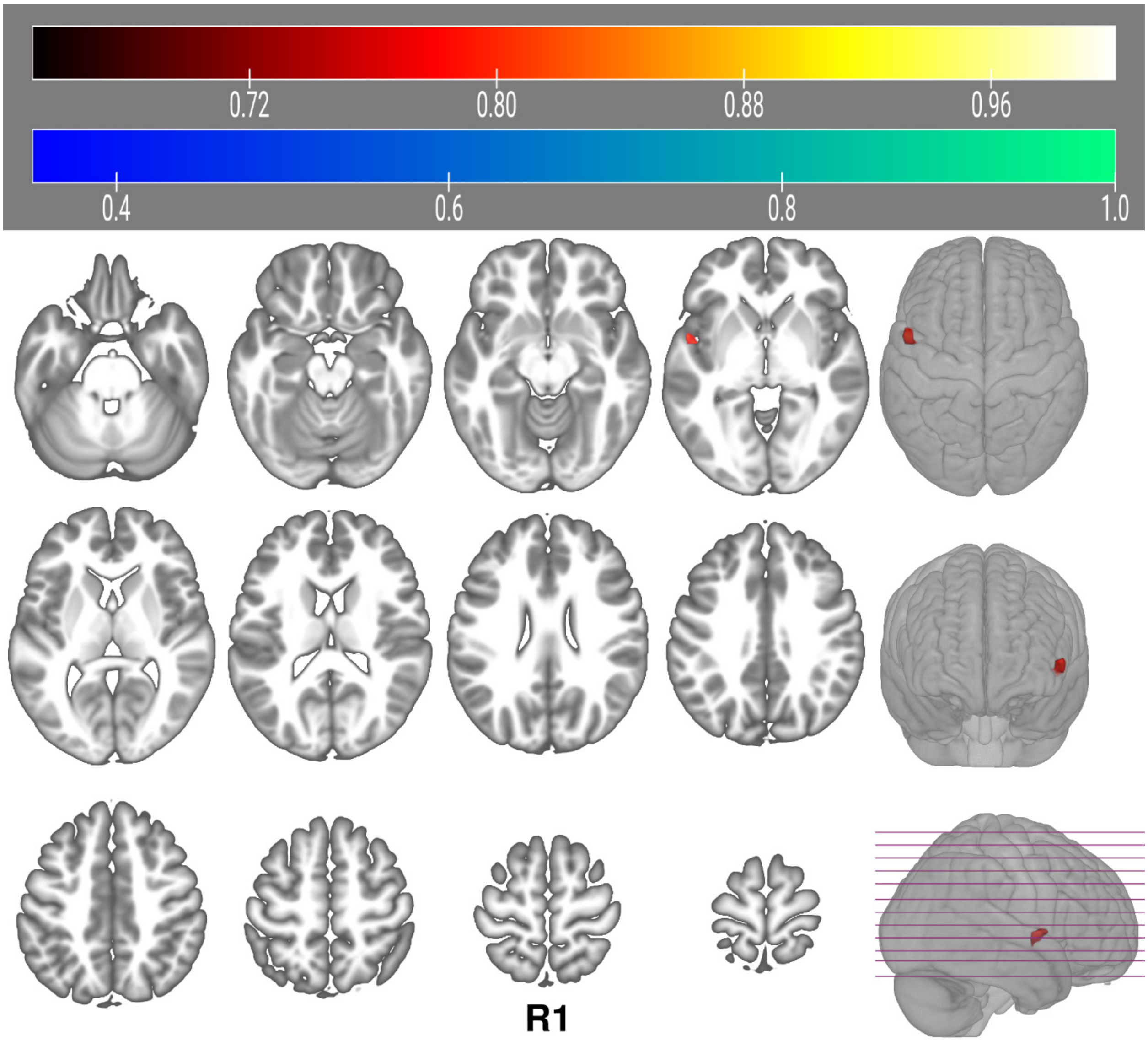

Figure S5.

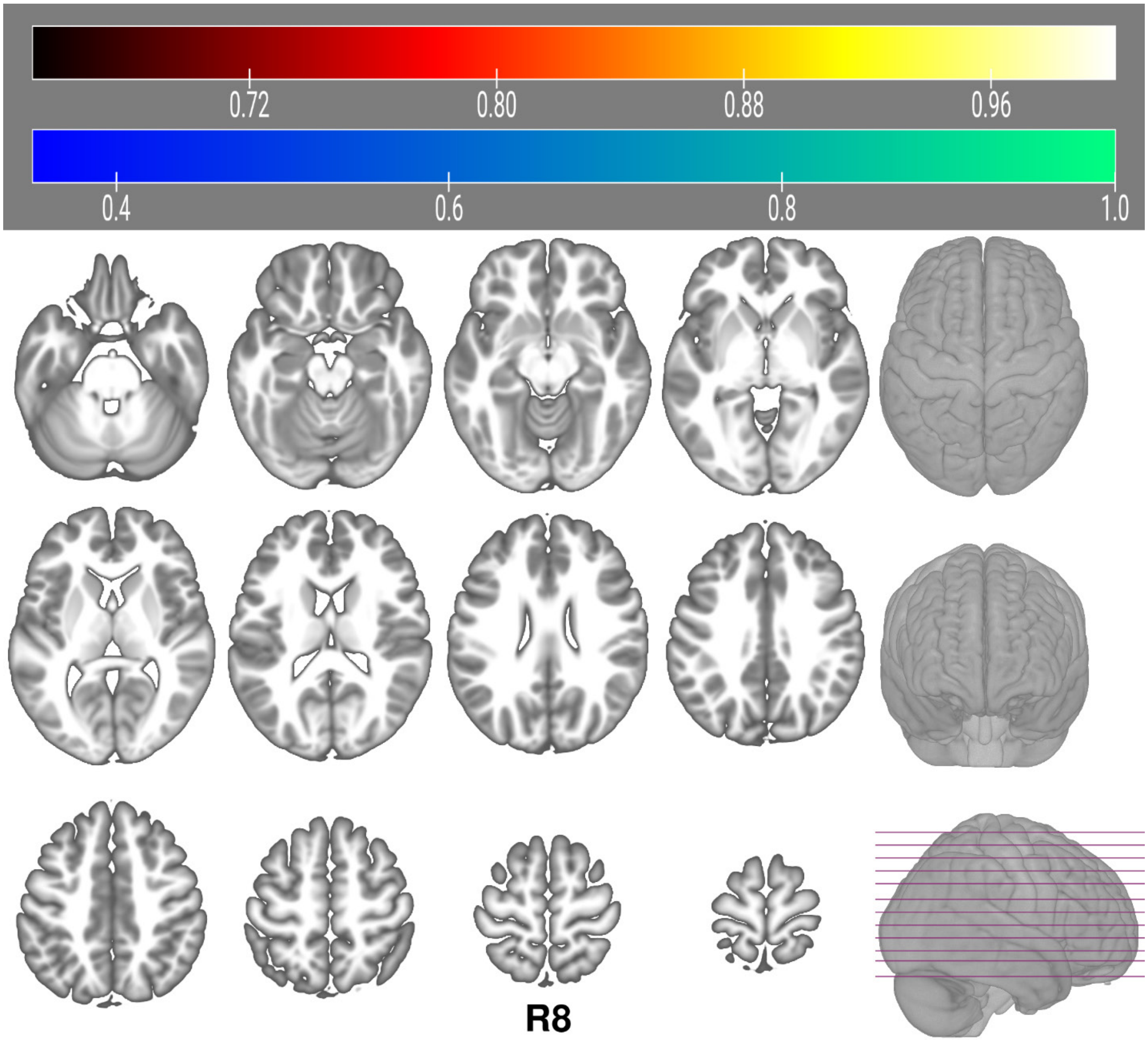

Dynamic multimodal components

Dynamic multimodal components (Figure S6-S10). It shows more differences between HC and SZ across two States for the top five dynamic components compared to top five stable components, in both structural and functional connectivity.

Figure S6.

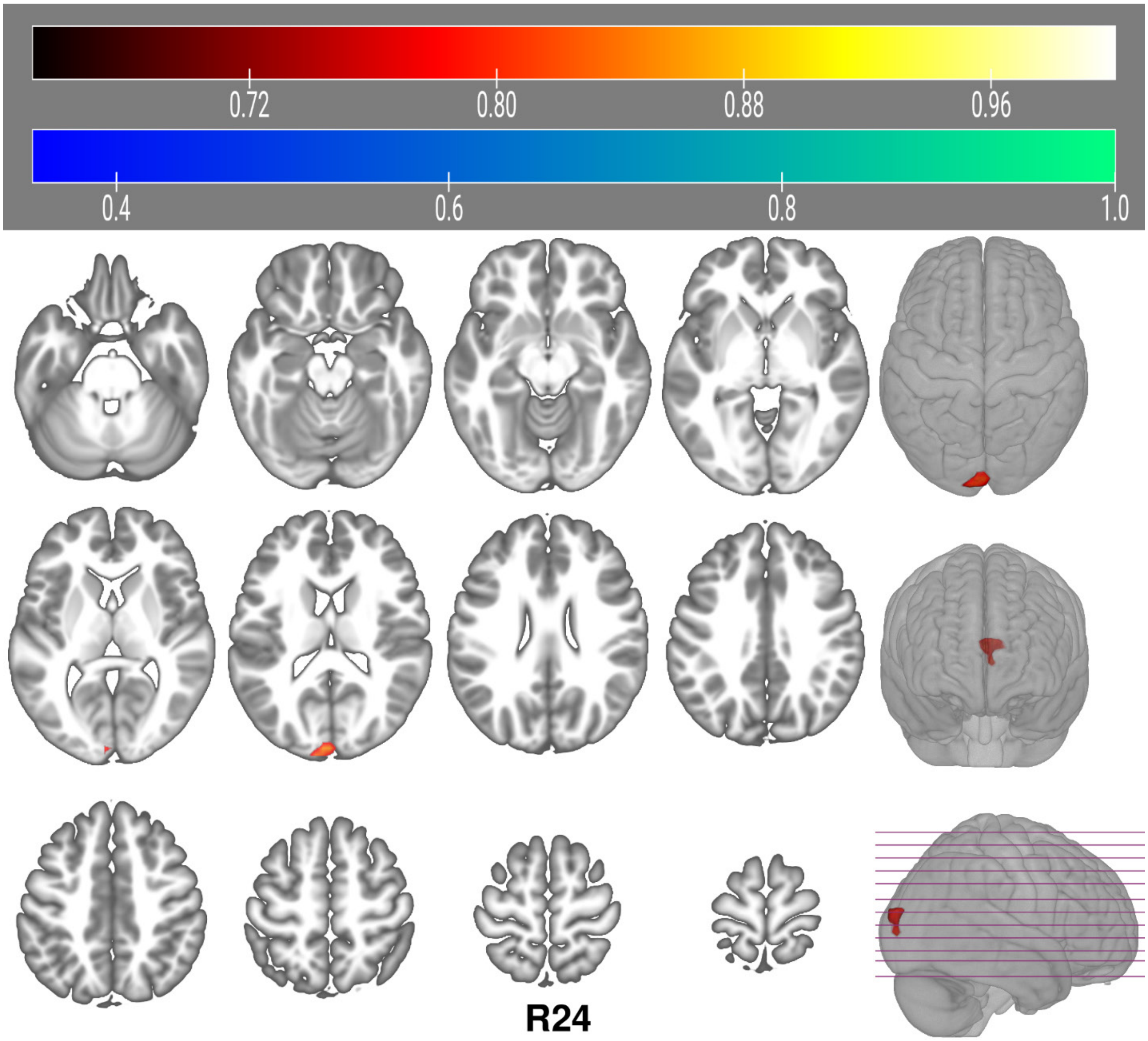

Figure S7.

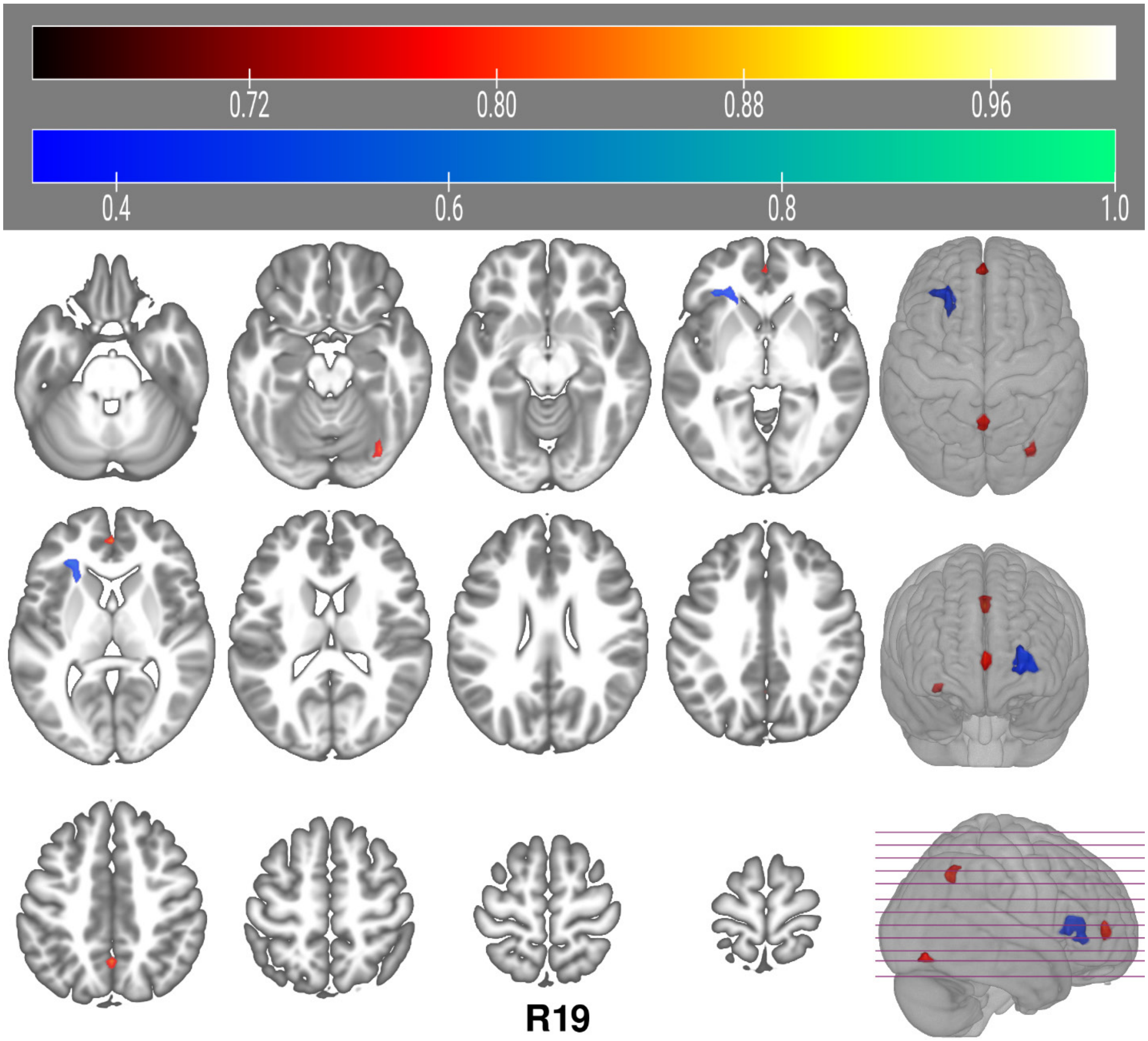

Figure S8.

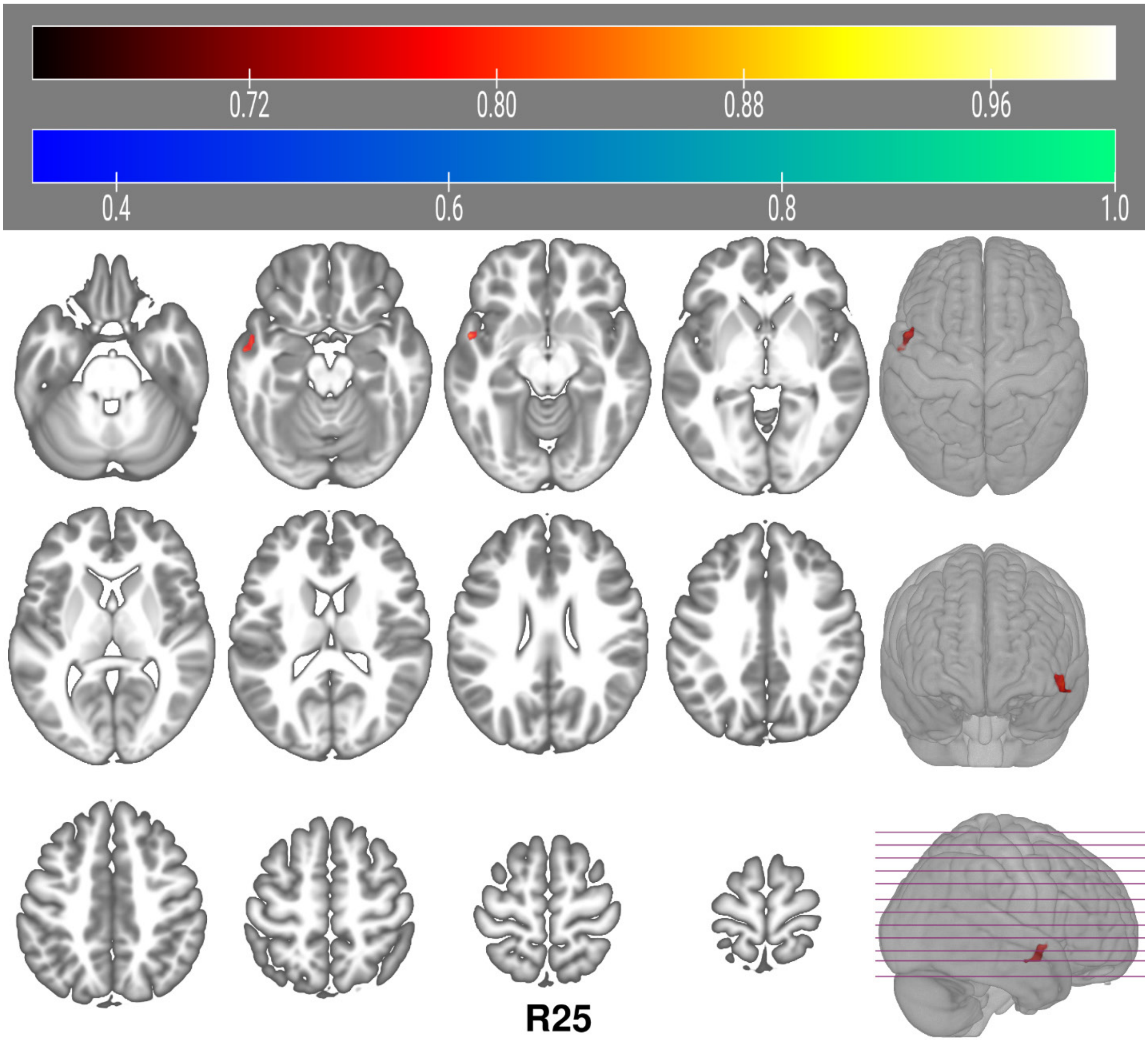

Figure S9.

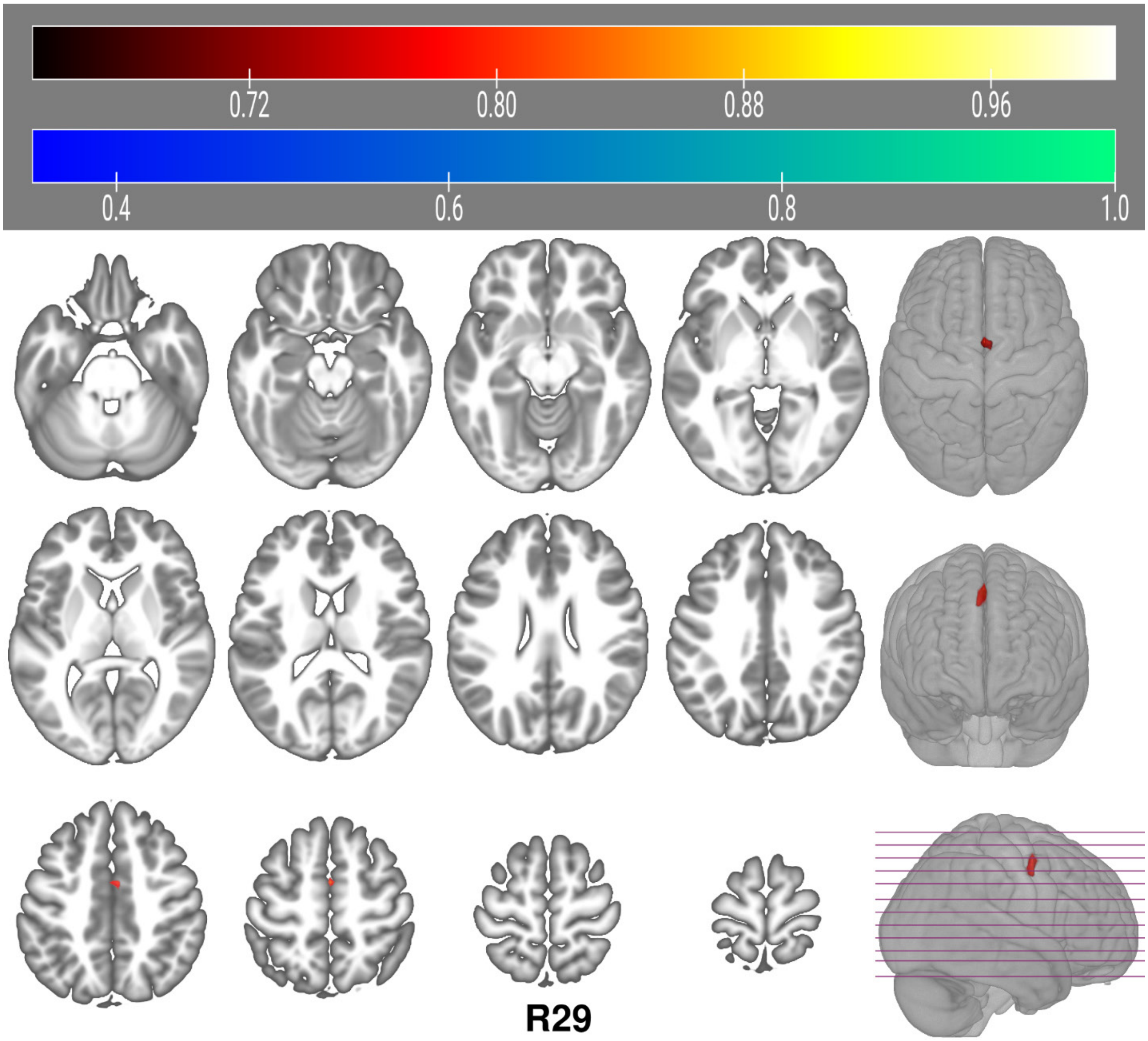

Figure S10.

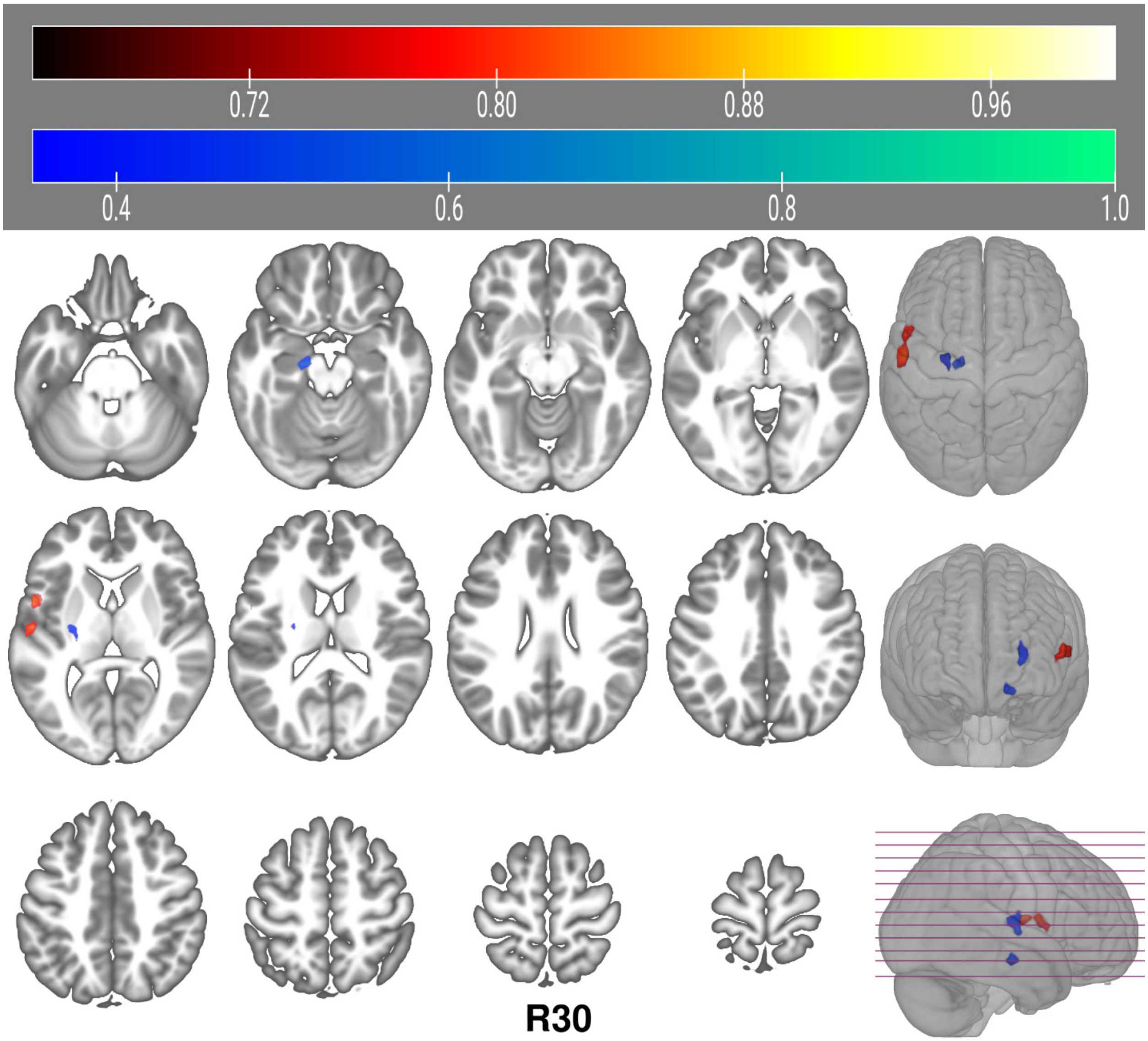
